# Disordered climate threatens short-distance migrants

**DOI:** 10.1101/2021.07.27.453912

**Authors:** Haile Yang, Luxian Yu, Hongfang Qi, Shengyun Fu, Yang Wang, Jianxin Yang, Hao Du

## Abstract

Global climate change has led to a warmer world, changing the migratory and breeding behaviors of many species, and short-distance migrants may benefit from climate change. With climate change leading to an increasingly disordered climate, we show here that a disordered spring climate disturbs the migration and breeding of a short-distance anadromous fish. In 2020, on the Qinghai-Tibetan Plateau, an abnormally low temperature in April delayed the migration rhythm of *Gymnocypris przewalskii* by nearly 10 days, while the gonadal development rhythm of the breeding population was almost normal. The phenology mismatch decreased the migrating populations by 30–70%, reducing the larval flux by nearly 80%. This case reveals that for short-distance migrants, different phenologies within the same species respond to disordered climates differently, which leads to phenology mismatches and then threatens the species. Along with increasing local extreme weather and climate events, short-distance migrants need more attention and conservation actions.

## INTRODUCTION

Climate change alters every biome and affects nearly every species (Freeman et al. 2018; Riddell et al. 2019; Barbarossa et al. 2021; Crozier et al. 2021), especially migratory species (Bowers et al. 2016; Wegge & Rolstad 2017; Tomotani et al. 2018; Cohen & Satterfield 2020; Paumier et al. 2020). Species with shorter migration distances are better able to predict the onset of spring at their breeding sites than species with longer migration distances do (Visser et al. 2009). Therefore, short-distance migrants appear to respond to changes in temperature, mid-distance migrants respond particularly to holistic climate drift, while long-distance migrants respond to climate change with a long time lag or even tend to not change over time (Miller-Rushing et al. 2008). The spring migration of long-distance migrants relies on endogenous rhythms that respond to climate change with a long time lag or is not affected by climate change (Both & Visser 2001), and then the populations and the food for provisioning nestling peaks are mismatched (Sanz et al. 2003; Both et al. 2006; Møller et al. 2008; Visser & Gienapp 2019; Zhemchuzhnikov et al. 2021). In contrast, along with the onset of spring advancement, as part of the environmental stress is released, the breeding success of short-distance migrants could be enhanced (Wegge & Rolstad 2017). Therefore, short-distance migrant populations have increased in response to climate change, while long-distance migrant populations have declined (Pearce-Higgins et al. 2015). However, would short-distance migrants always benefit from climate change?

Climate change not only involves increases in global average temperature and shifts in average precipitation but also leads to increases in the frequency and intensity of extreme weather and climate variability (IPCC 2013; Stott 2016; Swain et al. 2018). Individuals adjust their phenology along with environmental variables (such as photoperiod, temperature, rainfall and development of vegetation) due to their ectothermic physiology or use environmental variables that are predictive of the ‘optimal time window’ (McNamara et al. 2011; Visser & Gienapp 2019). Species differ in the relative importance of the different variables that affect their phenology and in the ways they respond to them (Visser & Gienapp 2019). As climate change has not led to uniform climate drift (such as temperature increase), even two species that rely on the same environmental variables have different phenology drifts (Gienapp et al. 2014; Renner & Zohner 2018). Moreover, the phenology of different annual cycle stages within the same species differs in response to the same climate change (Tomotani et al. 2018). Then, would the mismatch between different phenologies within the same species driven by a disordered climate be a path to threaten species?

Here, we proposed that a short-distance migratory species could be threatened by a disordered climate according to a phenology mismatch within the same species. To test this hypothesis, we focused on a short-distance anadromous fish (*Gymnocypris przewalskii*) to research whether and how a disordered climate impacts the breeding of this species. We monitored two phenology indicators, spring migration (arrival dates) and gonadal development (Ⅳ+), and two population dynamics indicators, migrating population size and larval flux. Following the framework of environmental drivers of variation and flexibility in fish migration (Tamario et al. 2019; Paumier et al. 2020), we monitored two environmental indicators: air temperature (indirectly indicating river water temperature) and river discharge. Furthermore, we analyzed the variation in spring migration, gonadal development, migrating population size, larval flux, air temperature and river discharge to verify our hypothesis. Moreover, we discussed the potential risk of the short-distance migrant fitness consequences due to disordered climate.

## MATERIALS AND METHODS

### Study species and study area

*G. przewalskii*, Qinghai Lake naked carp, a short-distance anadromous fish, is the predominant species in Qinghai Lake (Wang et al. 2013). In Qinghai Lake, *G. przewalskii* needs more than 7 years to reach sexual maturity, and then its breeding group seasonally migrates to inflowing rivers to spawn over sandy gravel beds when the river water temperature rises to 6~17.5°C from April to July (Xiong et al. 2010). The length of inflowing rivers is shorter than 290 km, and the migration distance of *G. przewalskii* is mainly 3 km to 120 km.

Qinghai Lake (N36.51°–37.25°; E99.58° –100.79°), the largest inland saltwater lake in China, is located on the Qinghai-Tibetan Plateau. The Qinghai-Tibetan Plateau, named the third pole, is one of the most sensitive regions to global warming on Earth (Zhang et al. 2018; Yao et al. 2019; You et al. 2019). The average annual air temperature on the plateau increased by 0.319 °C/10 y during 1987-2016, whereas the value was 0.415 °C/10 y from 2005-2016 (Tang et al. 2018). From 1979 to 2016, in Qinghai Lake, the freeze start date and freeze completion date were pushed back by 6.16 days and 2.27 days, respectively, while the ablation start date and ablation completion date advanced by 11.24 days and 14.09 days, respectively (Cai et al. 2017).

### Data collection

We monitored the arrival dates of *G. przewalskii* breeding population migrated into the main spawning grounds (the reach between the railway bridges and highway bridges) of three main inflowing rivers (Buha River, Quanji River and Shaliu River) during 2014-2020. We monitored the breeding population size and larval flux of *G. przewalskii* in the main spawning grounds of the three main inflowing rivers during 2018-2020. From 2018 to 2020, there were 7 surveys in Qinghai Lake near the Buha River estuary in which *G. przewalskii* was captured using gill nets, and each individual was dissected to identify its stage of gonadal development. We collected the discharge records of the three main inflowing rivers from 2018 to 2020. We collected air temperature records from Gangcha during 2011-2020.

### Data analysis

We compared the migration rhythm variation of *G. przewalskii* with the variations of air temperature and river discharge to identify whether and how environmental variations impacted the migration phenology of *G. przewalskii*. We compared the migration rhythm variation of *G. przewalskii* with the gonadal development rhythm variation of *G. przewalskii* to identify whether there was a mismatch between migration rhythm and gonadal development rhythm. We analyzed the breeding population size and larval flux of *G. przewalskii* in the main spawning grounds of the three main inflowing rivers to identify whether the species was threatened.

## RESULTS

### Migration time delay of *G. przewalskii* and its consequence

In 2020, we noted that the migration time of *G. przewalskii* was delayed by nearly 10 days (Table 1) and that the migrating population of *G. przewalskii* declined by 30–70% in the three main inflowing rivers: the Buha River (nearly 30%), Quanji River (nearly 60%) and Shaliu River (nearly 70%). Subsequently, the larval flux of *G. przewalskii* that migrated into Qinghai Lake declined by nearly 80% (Table 1).

**Table 1.** The migration time and larval flux of *G. przewalskii* in three rivers

| Year | Time of *G. przewalskii* population migration into the lower boundary of the traditional spawning reach of each river | | | Larval flux of *G. przewalskii* migration from each river into Qinghai Lake ($\times 10^8$) | | |
| --- | --- | --- | --- | --- | --- | --- |
|  | Buha | Quanji | Shaliu | Buha | Quanji | Shaliu |
| 2014 |  | 4-Jun | 4-Jun |  |  |  |
| 2015 |  | 2-Jun | 2-Jun |  |  |  |
| 2016 |  | 2-Jun | 2-Jun |  |  |  |
| 2017 |  | 6-Jun | 6-Jun |  |  |  |
| 2018 |  | 29-May | 3-Jun | 3.91 | 0.37 | 0.38 |
| 2019 | 4-Jun | 3-Jun | 3-Jun | 2.82 | 0.16 | 0.86 |
| 2020 | 14-Jun | 11-Jun | 11-Jun | 0.44 | 0.09 | 0.06 |

### River discharge was not the factor that disturbed *G. przewalskii* migration in 2020

Considering that the starting time of *G. przewalskii* migration into the inflowing rivers was mainly in May and June (Table 1), we compared the discharge records of the three main inflowing rivers (Buha River, Quanji River, Shaliu River) in April, May and June from 2018 to 2020. The results showed that there was no obvious discharge signal difference between 2020 and 2018 or 2019 that could match the delay of *G. przewalskii* migration in 2020 (Figure 1). In other words, we inferred that river discharge was not the factor that disturbed *G. przewalskii* migration in 2020.

**Figure 1.**
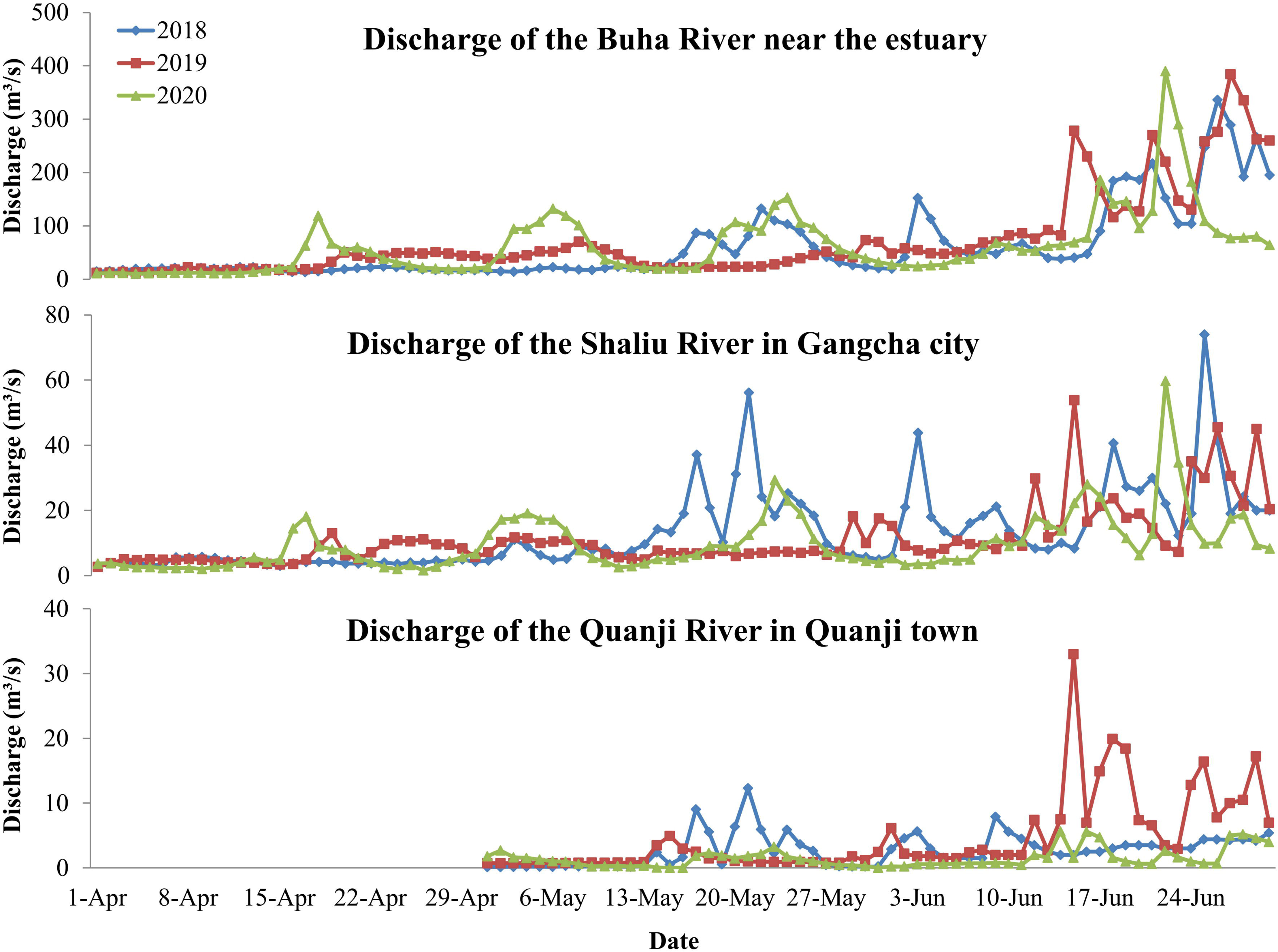
The discharges of the three main inflowing rivers. During April, the discharge of the Quanji River in Quanji town was too low to record.

### Delay of warming in April caused the delay of *G. przewalskii* migration in 2020

Considering that the starting time of *G. przewalskii* migration into the inflowing rivers was mainly in May and June (Table 1) and that there was a time lag between river water temperature and air temperature, we compared the air temperature records from Gangcha in March, April and May from 2011 to 2020. The results showed that the daily maximum temperatures in April were obviously lower in 2020 than in other years, and there was no obvious difference in March and May between 2020 and other years (Figure 2). In other words, there was a delay of warming in April 2020. Considering the time lag between river water temperature and air temperature, the delay of warming in April 2020 caused the delay of the river water temperature signal inducing *G. przewalskii* migration and then caused the delay of *G. przewalskii* migration in 2020.

**Figure 2.**
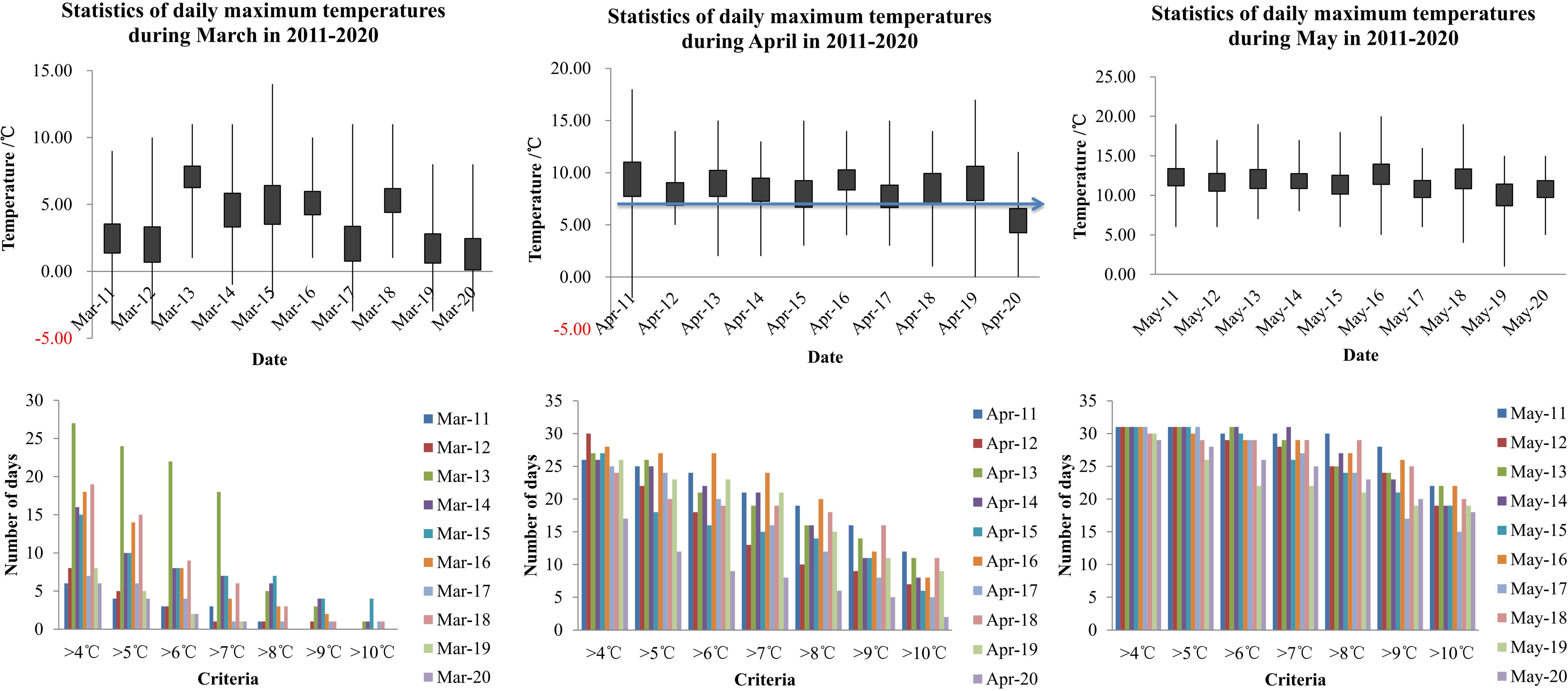
Statistics of daily maximum air temperature in Gangcha

### Phenology mismatch between the migration rhythm and gonadal development rhythm

We analyzed records on the gonadal development stage of each *G. przewalskii* individual captured in Qinghai Lake near the estuary from 2018 to 2020. The ratio of gonadal development (IV+) of the *G. przewalskii* population in each survey was calculated. The results showed that 19.35% of *G. przewalskii* individuals captured in Qinghai Lake near the Buha River estuary in the survey on 5 June 2020 were identified as postspawning. Compared with 2018 and 2019, there was no obvious delay in the gonadal development of *G. przewalskii*, and the percentage of individuals with gonadal development (IV+) was no obvious decline (Figure 3). Compared with the migration time delay, there was a phenology mismatch between the migration rhythm and gonadal development rhythm.

**Figure 3.**
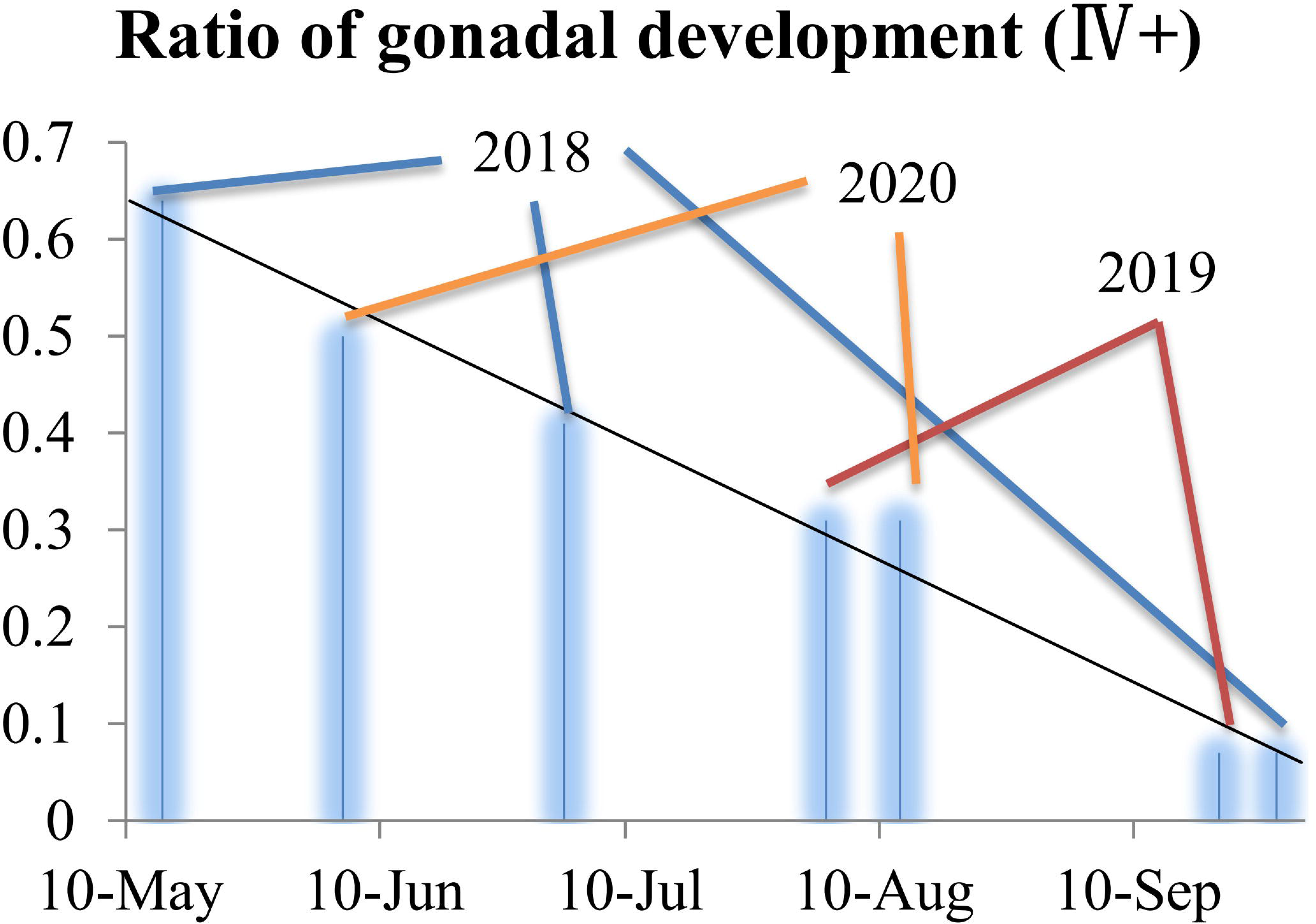
The ratio of gonadal development (IV+) of the G. przewalskii population in 7 surveys

## DISCUSSION

Phenology, the seasonal timing of life-cycle events, exists during which environmental conditions are most advantageous, i.e., an ‘optimal time window’ (Visser & Gienapp 2019). Many species shift their phenology in response to global climate change but often do not shift at the same rate (Kharouba et al. 2018). In particular, for migratory species, short-distance migrants are more sensitive to climate change than long-distance migrants (Miller-Rushing et al. 2008). Therefore, under climate change, short-distance migrants have a higher fitness than long-distance migrants, and then populations increase (Pearce-Higgins et al. 2015). However, climate change involves both the long-term change in temperature and precipitation as well as short-term disturbances in weather and climate (IPCC 2013; Stott 2016; Swain et al. 2018), which could lead to a mismatch between different phenologies within the same species (Tomotani et al. 2018). We proposed that short-distance migrants may be impacted by phenology mismatches within species that are driven by disordered climates and do not always benefit from climate change. In the present work, we showed that driven by the abnormally low (daily maximum) air temperature in April, the migration time of *G. przewalskii* (a short-distance anadromous fish) was delayed by nearly 10 days, which led to the phenology mismatch between migration and spawning; then, the migrating population declined by 30–70%, and the larval flux declined by nearly 80%. This case indicates that a disordered climate could cause a phenology mismatch within a short-distance migrant and then threaten the species.

It is clear that in response to climate change, different species shift their phenology at different rates, which causes mismatches between the phenology of interacting species and then leads to a series of evolutionary and population consequences (Renner & Zohner 2018; Samplonius & Both 2019; Visser & Gienapp 2019). In response to climate change, the phenology of different annual cycle stages within the same species also shift at different rates (Tomotani et al. 2018). Different phenologies respond to climate change with different sensitivities. Sensitive phenology always drifts with short-term environmental indicators, such as temperature. Stabilized phenology always drifts with long-term climate indicators, such as the effective accumulated temperature. If the two phenologies of a species shift at different rates along with different environmental indicators, there would be a phenology mismatch. In the present work, the gonadal development rhythm of *G. przewalskii* responded to climate change with low sensitivity, and the migration rhythm of *G. przewalskii* responded to climate change with high sensitivity. From 1979 to 2016, in Qinghai Lake, the freeze start date and freeze completion date were pushed back by 6.16 days and 2.27 days, respectively, while the ablation start date and ablation completion date advanced by 11.24 days and 14.09 days, respectively (Cai et al. 2017). The average annual air temperature on the plateau increased by 0.319 °C/10 y during 1987-2016, whereas the value was 0.415 °C/10 y from 2005-2016 (Tang et al. 2018). We infer that the gonadal development rhythm of *G. przewalskii* has advanced gradually in past decades, although there may be a time lag. Based on the survey results on the gonadal development of *G. przewalskii* in 2018, 2019 and 2020 (Figure 3) and considering the historical records from the 1970s to 1990s that *G. przewalskii* migrates to inflowing rivers to spawn from April to July (Xiong et al. 2010), we identify that the breeding time window opens in April. In contrast, driven by the abnormally low (daily maximum) air temperature (indirectly indicating river water temperature) in April 2020, the migration rhythm of *G. przewalskii* was delayed by nearly 10 days. The mismatch between the migration window and the breeding time window caused part of the *G. przewalskii* breeding group to spawn outside the traditional spawning habitats in 2020. Then, the breeding population that migrated into the traditional spawning habitats and the larval flux that migrated from the spawning habitats into Qinghai Lake seriously declined.

With climate change leading to an increasingly disordered climate (IPCC 2013; Stott 2016; Swain et al. 2018), short-distance migratory species need more attention and conservation actions. As a disordered climate could severely impact sensitive phenology and stabilized phenology responds to climate change with a time lag, the mismatch between sensitive phenology and stabilized phenology within the same short-distance migrants would impact their breeding success, although phenotypic plasticity could provide the potential for organisms to respond rapidly and effectively to environmental change (Paumier et al. 2020; Senner et al. 2020). In our survey conducted on 5 June 2020, 19.35% of *G. przewalskii* individuals captured in Qinghai Lake near the Buha River estuary were identified as postspawning when the *G. przewalskii* breeding group had not migrated into the traditional spawning habitats. In other words, some fish spawned in the estuary or adjacent bay. Perhaps spawning in these other areas is a possible way for *G. przewalskii* to adapt to climate change. However, whether the breeding is successful and whether there are enough healthy larvae and juveniles in the new nonriver spawning sites need to be investigated because this determines the population dynamics of *G. przewalskii*. Disturbed by extreme events and then spawning at a substitute site, the attempt almost failed in the story of Chinese sturgeon (Huang & Wang 2018). Therefore, we need more attention and conservation actions for *G. przewalskii* and other short-distance migrants and hope there could be positive outcomes.

Perhaps different taxa (such as mammals, birds and fish) of migrants have different responses to climate change. We believe that the processes by which climate change impacts migrants are general. In the present work, we identified that a disordered climate could sensitively disturb the migration of short-distance migrants, while gonadal development and breeding rhythms were stabilized. The phenology mismatch led to the decline of breeding populations in traditional spawning locations and the decline of newborn offspring. For fish, spawning adults and embryos are the most critical life stages and are very sensitive to temperature (Dahlke et al. 2020). Therefore, the abnormal cold April delayed the migration of *G. przewalskii* in 2020 and then caused a series of serious consequences. Other taxa (such as mammals and birds) also have critical bottlenecks in their life cycle. A corresponding disordered climate (or environmental) variability could also lead to a series of serious consequences. Migrants may adapt to gradual climatic shifts through phenotypic plasticity and even evolutionary adaptability (Bowers et al. 2016; Senner et al. 2020). However, abnormal and abrupt climate change would be dangerous.

In summary, using a case study on a short-distance anadromous fish, we verified that a disordered climate could disturb the sensitive phenology of short-distance migrants but did not impact their stabilized phenology, which causes phenology mismatch within the same species and then threatens the species. Whether migrants could adapt to this abnormal and abrupt climate change is unknown. Following increasingly extreme weather and climate variability, we need more attention and conservation actions for short-distance migrants.

## Acknowledgements

This work was supported by the Natural Science Foundation of Qinghai [grant numbers 2018-ZJ-908]; and the National Freshwater Fisheries Germplasm Resources Conservation Bank Project [grant numbers FGRC-18537].

